# Understanding the effects of moisture content and temperature on dormancy release in sunflower (*Helianthus annuus* L) achenes

**DOI:** 10.1101/2022.12.22.521709

**Authors:** Gonzalo Joaquín Arata, Diego Batlla, Patricia Verónica Demkura, María Verónica Rodríguez

## Abstract

The effects of moisture content (MC) and storage temperature (ST) on seed longevity have been modeled for many species. In contrast, our understanding on the combined effects of MC and ST on dormancy release (DR) in “dry” orthodox seeds is still insufficient to build robust predictive models. We used freshly harvested, dormant sunflower achenes to explore the effects of MC (4-10%) in combination with a wide range of ST (−18°C to +30°C) on DR dynamics, embryo responsiveness to abscisic acid (ABA) and deterioration indicators. Storage temperatures allowing full DR were inversely related to achene MC, ranging from >25°C for MC4% to sub-zero temperatures for MC10%, resembling a phase diagram. Rates of DR were plotted along a RH gradient. Combinations of MCxST optimal for DR were between *ca*. 40-60%RH. Increasing RH from 60 to 80% inhibited DR. Higher RH>80-85% promoted partial DR together with rapid ageing. We suggest that reactions promoting full DR are favored alongside a physical (glassy) transition and are not oxidative. We propose biophysical coordinates to guide future studies on the mechanisms involved in DR, but also to develop predictive models useful to define post-harvest conditions that maximize sunflower seed quality.

**Highlights:**

□ The effects of moisture content (MC) and storage temperature (ST) on dormancy release and deterioration were investigated in sunflower achenes using a factorial design.
□ Dormancy release was promoted by MCxST combinations in equilibrium with a RH between 40 and 60% and was delayed outside this range.
□ Storage temperatures optimal for dormancy release were inversely related to achene MC, ranging from >25°C for MC4% to sub-zero temperatures for MC10%, resembling a phase diagram.
□ Dormancy release and ageing are promoted within distinct, non-overlapping regions along the RH gradient supporting different types of reactions for each process.

## Introduction

Seed dormancy is an internal block for germination with a clear adaptive value in wild species (Finch-Savage and Leubner-Metzger, 2006). However, in crop species prolonged dormancy can interfere with proper crop establishment (Benech-Arnold *et al*., 2012). This is usually a problem during sunflower hybrid “seed” (strictly, a fruit-achene or cypselae-) production, in which achenes are dormant at harvest and require a period of dry after-ripening (AR) to become non-dormant and meet germination standards, i.e., high germination rates at both 10 and 25°C (ISTA 1999). This period may last from a few weeks to several months or over a year depending on the genotype (Arata *et al*., 2021) and is influenced by the maternal (Bodrone *et al*., 2017; Lachabrouilli *et al*., 2021; Riveira Rubin *et al*., 2021) and post-harvest environments (Rodríguez *et al*., 2018; Bazin *et al*., 2011a). Although domestication and breeding have reduced primary dormancy in many crops (as in the case of cereals), among cultivated sunflower germplasm many elite lines still display dormancy attributes comparable to their wild relatives (Arata *et al*., 2021).

Orthodox seeds are characterized by a low moisture content (MC) in equilibrium with air humidity at dispersal/harvest maturity (Baskin and Baskin, 2004). In this very low hydration range, the cytoplasm enters a solid or glassy state at ambient temperatures that restricts molecular mobility (Leopold *et al*., 1994; Ballesteros and Walters, 2011, 2019). Reactions in the glassy state are limited to solid state oxidation, peroxidation, and carbonylation of molecules in proximity within the solid cytosol (Ballesteros and Walters, 2019). As these reactions lead to cumulative damage of seed components, seeds age and become non-viable. Reactive oxygen species (ROS) might also have a role in dormancy release, through oxidation of mRNA and proteins in a specific (regulatory?) way (Oracz *et al*., 2007; Bazin *et al*., 2011b; Meimoun *et al*., 2014; Bailly *et al*., 2019). Nevertheless, a causal link between endogenous ROS formation in the dry seed and dormancy release has not been fully established yet, and the existence of a functional relationship (if any) between dormancy release and oxidative reactions also involved in seed ageing is still unclear.

The combined effects of moisture content and temperature during storage on seed ageing dynamics have been the subject of many thorough and detailed studies that allowed the development of empirical and robust seed longevity models (Ellis and Roberts, 1980; Cromarty *et al*., 1982). Seed MC depends on air relative humidity (RH), temperature and seed composition (Pixton and Warburton, 1973). The RH is used as an estimate of seed a_*w*_ and can be related broadly to the seed’s metabolic activity (Vertucci and Leopold, 1986; Leopold and Vertucci, 1989). The relationship between MC and RH is described by sorption isotherms, which also provide information on the binding properties of water and the type of reactions that prevail along the RH gradient. Vertucci and Roos (1990, 1993) proposed the use of RH to define storage protocols to maintain seed longevity. An updated discussion on their importance in longevity and dormancy studies was recently published by Hay *et al*. (2022).

Contrary to studies on seed longevity, those focused on the effects of the storage environment on dormancy release are comparatively limited. A main concern about these studies is that methodologies employed to manipulate MC are diverse and may affect the results and their interpretation. For example, when using saturated salt solutions to allow seeds to equilibrate with a target RH, time to reach a final MC can last several weeks, long enough to overlap temporally with changes in dormancy status. Another constraint is the narrow (or incomplete) range of storage conditions that were tested within the same study. For example, in sunflower, Bazin *et al*. (2011a) studied the combined effects of MC and storage temperature (ST) on dormancy alleviation in achenes with MC between 3 and 12% but, within this range, they omitted MC between 5 and 10 % which are commonly reached under “ambient” conditions (i.e., RH 40-80%; Pixton and Warburton, 1971). Also in sunflower, Rodríguez *et al*. (2018) observed a promoting effect of warmer ST for achenes dried to MC 6% but stored non-hermetically, so MC may have changed during storage at different temperatures. A better description of how MC and ST influence dormancy release is needed to solve a practical agronomic problem (i.e., post-harvest conditions for sunflower hybrid seed production) but also to gain more insight into the mechanisms involved in the dry after-ripening process in this and other species as well.

The effects of MC and temperature on dormancy release are far from simple, and significant interactions can be expected. Very often, higher AR temperatures accelerate DR (Roberts, 1965; Baskin and Baskin, 1976, 1986; Allen *et al*., 1995; Probert RJ., 2000). However, the response to AR temperature in some studies depended on MC. An inverse relationship between temperatures promoting DR and seed MC was reported in wild oat (Foley, 1994), while the opposite pattern was reported in *Arabidopsis* (Basbouss-Serhal *et al*., 2016) and sunflower (Bazin *et al*., 2011a). Nevertheless, these results are not directly comparable due to methodological differences and ranges of MC and ST used in each study (see Supplementary Table S3).

Whatever the subtle changes are that take place during dry storage, they have a strong impact on the physiology of the imbibed seed and its germination response (Chahtane *et al*. 2017). In Arabidopsis and barley, imbibed, dormant seeds exhibit differences in ABA metabolism and signaling as compared to imbibed, after-ripened (non-dormant) seeds (Ali-Rachedi *et al*., 2004; Benech-Arnold *et al*., 2006). In sunflower, ABA synthesis maintains embryo dormancy during achene development and maturation (Le Page-Degivry *et al*., 1990, 1996; Le Page-Degivry and Garello, 1992). However, ABA metabolism during imbibition is not clearly related with dormancy alleviation of sunflower achenes as they after-ripen (Rodriguez *et al*., 2018). In parallel, sensitivity to ABA, assessed as the inhibition of embryo germination by exogenous ABA, progressively decreased during dry storage (Bianco *et al*., 1994; Le Page-Degivry *et al*., 1996), and was associated with the different dormancy release dynamics obtained at two storage temperatures (5 vs. 25°C; Rodriguez *et al*., 2018). The cause of this decreased responsiveness to ABA is unknown but may involve direct changes in the functionality of components of the ABA signaling pathway, or other hormonal pathways known to antagonize the ABA signal, such as gibberellins and ethylene (Corbineau *et al*., 1990, 2014; Corbineau and Côme, 2003; Xia *et al*., 2019).

The aims of this work were to investigate the effects of storage temperature and MC on DR of sunflower achenes over a broad range of RH conditions, integrating physiological information for dormancy and deterioration. Finally, we discuss if both dormancy release and seed ageing have a common mechanistic basis or are regulated by distinct, independent mechanisms.

## Materials and methods

### Plant material

Field trials were conducted in the experimental field at the Facultad de Agronomía de la Universidad de Buenos Aires (FAUBA) (34°35’37”S 58°29’03”O) during spring and summer in 2017-2018, 2018-2019 and 2019-2020 and are referred to as experiments 1, 2 and 3, respectively. Field plots were always sown within October. Experiments were performed with publicly available inbred lines: “600” (source: developed by the Instituto Nacional de Tecnología Agropecuaria and provided by INTA Pergamino; derived from Russian × USDA North Dakota, oil content 49%); “1579” (USDA North Dakota, oil content 40%). Both “600” and “1579” were used in *experiment 1*, while experiments 2 and 3 were conducted only with inbred line “600”. According to previous work (Arata *et al*., 2021), dormancy phenotype of “600” is intermediate-low with no thermo-inhibition, while “1579” displays deeper dormancy and thermo-inhibition when incubated at 30°C. Field experiments were irrigated and fertilized, and pest control was carried out following standard practices. Plant density was 5 plants/m^2^ and distance between rows was 52 cm. Cross-pollination between lines was avoided by covering heads in R4 stage with medium-heavy polyester bags. Pollination was facilitated manually with a soft paint brush. At harvest maturity (*ca*. 11% MC), 20-30 heads of healthy and phenologically synchronous plants were harvested and threshed manually. Achenes from the center and the two outermost peripheral rows were discarded. The rest of the fruits were pooled together to obtain a main pool.

### Storage treatments

Each lot of freshly harvested and threshed achenes was separated into four sub-lots *(ca*. 500-600 g), each one destined to reach a target moisture content (MC 4, 6, 8, 10%, dry weight basis). Moisture content was determined gravimetrically (ISTA, 1999). Two different procedures were followed to obtain the target MC. In *experiment 1* (season 2017-2018) the four sub-lots were dried to 6% using an experimental dryer (air flow at 35°C). Two of these sub-lots were then placed in a saturated atmosphere (100% RH) for 21 d at 20°C to reach MC 8 and 10%. Increase in MC was monitored twice a day by weighting each sample. A remaining sub-lot with MC 6% was placed in a container with silica gel (1:1 w/w, replenished twice per day) at 20°C until MC reached 4%. In the following experiments (2 and 3) a desorption schedule was followed immediately after harvest. Sub-lots were placed in the experimental dryer (air flow at 35°C) and removed sequentially when each target MC was reached (10, 8 and 6%). The lowest MC (4%) was obtained by drying over silica as described above. Temperature during drying over silica was kept at 20°C, as well as the other sub-lots after reaching their respective target MC (6, 8 and 10%) until initiation of storage treatments of all samples together.

Time to reach the target MC was about 21 d for *experiment 1*, and this was shortened to 7-9 d in *experiments 2* and *3* in which all samples were obtained by desorption (the last sample to reach target MC was 4%). Once the four target MC had been reached, 10 g aliquots of achenes (ca 150 achenes) were placed in glass containers (100 cm^3^ bottles) and closed with rubber stoppers hermetically sealed with vacuum grease and parafilm. For each MC, triplicate containers (n=3) were stored in chambers set at a constant temperature following a factorial design (4 MC × 6-7 storage temperatures). Storage temperatures always ranged from +5 to 25°C (with 5°C increments) and in some cases, −18°C (freezer) and +30°C. A separate set of containers was used for each sampling time (e.g. 30 and 70 d), so these remained closed until the germination assays. Additional storage treatments were included to investigate the effect of ambient oxygen. Anoxia was combined with MC 6 and 8% in *experiment 1*, and with all MC levels in *experiment 2*. Oxygen was replaced by a constant flow of N_2_ gas through two (in/out) needles inserted through the rubber stopper during 1 min. After removing the needles, surface of the rubber stopper was sealed with nail polish, and excess pressure was released before it solidified. These treatments were stored at +5 and 25°C. The anoxic condition was confirmed again at the end of storage with an O_2_ needle-type sensor (Presens, Neurburg, Germany).

### Monitoring of achene MC during storage

Moisture content (MC) was determined gravimetrically for all treatments and replicate samples (bottles) before and after each storage period. Ten achenes were weighed before (fresh weight, FW) and after (dry weight, DW) oven drying at 130°C for 120 minutes (ISTA, 1999). Moisture content values (%) are expressed on a dry weight basis.

### Germination tests

Dormancy level was assessed by incubating achenes and embryos at 10°C (which favors the expression of dormancy) and at 25°C (in experiments 2 and 3) or 30°C *(experiment 1)*, where dormancy is less expressed (see Arata *et al*., 2021). Incubating at two contrasting temperatures allows to distinguish among deep-dormant seed samples (e.g., soon after harvest, where differences will be best observed at 25°C) and among less-dormant samples where differences will be best observed at 10°C. Incubation at 30°C also allows to detect thermo-inhibition (30°C) depending on the genotype (e.g. “1579”; see Arata *et al*., 2021). Occasionally, imbibition was also performed in a solution of 200 uM Etephon (an ethylene donor) to promote germination of dormant achenes.

On each sampling date, germination assays were conducted in triplicate Petri dishes for each storage condition. Twenty-five achenes (or 20 embryos) per replicate were placed on top of two layers of laboratory filter paper and 6 ml of distilled water and incubated for 15 d in temperature-controlled chambers, with no additional light (but ambient light was not avoided during counts). Germinated units were scored every 2-3 d and removed. Achenes were considered to have germinated when the radicle protruded visibly 2-3 mm through the open pericarp. For embryos, germination was considered complete when the radicle had elongated >5 mm and began to curve. Germination assays were performed before initiation of storage (for the four MC treatments) and at least twice during storage. Sampling times were: 30/37 d, 70/77 d (for lines “600” and “1579, respectively”) and 200 d (only line “600”) in *experiment 1;* 30 and 70 d in *experiment 2;* and 30, 70 and 100 d in *experiment 3*.

In parallel to the incubation of embryos in water, embryos were incubated in 5 μM ABA (at 10°C in *experiment 1*, and 25°C in *experiment 2*).

### Oxygen measurements

The oxygen level in the headspace of glass bottles containing achenes was measured by inserting a needle-type micro-sensor through the rubber stopper (50 μm tips, Presens, Neurburg, Germany; methodology according to (Rolletschek *et al*., 2009; Dominguez *et al*., 2019). Ambient temperature was kept stable at 25°C during measurements. Data, obtained in mV, was converted to O_2_ concentration using a calibration curve (interpolated from known O_2_ concentration in ambient air and anoxia). These values were then expressed as relative to ambient O_2_ concentration. Measurements were done twice (*ca*. 30 and 70 d) during storage and for all storage treatments in *experiment 1*.

### Lipid peroxidation

Lipid peroxidation in the embryo axis was assessed by TBARS in samples stored for 70 d from *experiment 2*. For each replicate container (bottle), 15 achenes were used for TBARS. After removing the pericarp, 15 embryo axes were isolated with a scalpel and immediately ground in a mortar with sterilized sand, extracted in 20% trichloroacetic acid (TCA) and reacted in 0.5% TBA to 20% TCA at 95°C (Heath and Parker, 1968). A reaction blank with ground sand was included. The TBA reactive substances were determined spectrophotometrically at 532 nm and non-specific absorbance was determined at “600” nm. Results are expressed as MDA (mg) per g of embryo axis (fresh wight).

### Electrical conductivity

Twenty-five de-hulled seeds (only the pericarp was removed, without damageing the seed coat) were placed in 37.5 ml of de-ionized water (< 2 μS.cm^-1^ g^-1^) at 25°C as proposed by Szemruck *et al*. (2015). The electrical conductivity of the solution was measured after 24 h (at 25°C) using an Accumet AP85 portable conduct-meter, and results expressed in μS.cm^−1^ g^−1^ (adapted from Braz *et al*., 2008). Determinations were made at 70 DAS in genotype “600” of *experiment 1*, and EC was measured in each of the 3 replicates of each storage treatment. Data is presented as mean +/- SD.

### Quantification of endogenous ABA

Abscisic acid was measured by radioimmunoassay after a 70 d storage period in some treatments from *experiment 2*. These included MC 4, 6, 8 and 10% in combination with ST 5, 15 and 25°C. After 70 d of storage, 20 achenes were sampled from each replicate container. Pericarp was removed and the axis separated with a scalpel and stored at −80°C until processing. The samples were freeze-dried, ground to powder and used for aqueous extraction of ABA as described in Steinbach *et al*. (1995).

### Prolonged storage experiment

After *ca*. 200 d of storage, remaining achenes from “600” in *experiment 1* were used to test germination and seedling vigor. Achenes were incubated at 10°C and final germination scored on day 14 when it had reached a plateau. Achenes that did not germinate were transferred to Petri dishes with 6 ml of 100 μM ethephon (an ethylene donor) solution at 10°C, and germination was evaluated after four days. Germinated achenes were removed and placed in plastic trays (10×15×4cm) on top of a cotton layer covered with filter paper and soaked with distilled water, and cultivated in a growth chamber (22-23°C, 16 h photoperiod) for four days (SupplementaryFig. S 12 A). On day 4, seedlings were weighed individually. Data is presented for each storage treatment as the average FW of all seedlings ± SD (n = 20 to 25 seedlings).

### Moisture content shifting experiments

Freshly harvested achenes of genotype “600” (same original batch from *experiment 3)* were dried to reach MC 6, 8 and 10% (experimental air dryer at 35°C) and stored at 20°C. After 28 d of storage, a sub-sample (*ca* 40 g) from each MC condition was taken to modify its MC. Moisture content shifting was from 6 to 10% (11 d in humid chamber RH 100%), from 8 to 6% and from 10 to 6% (both drying under air flow at 35°C; 2-6 h). Achene germination was tested at 10 and 25°C 28 DAS (just before MC shifting began), 39 DAS (after MC shifting was complete for all treatments) and at 60 and 100 DAS. It should be noted that lowering MC by drying under air flow was faster than rising MC in a humid chamber; therefore, achenes dried to MC 6% “after-ripened” for 9 d with a fixed MC 6%, while MC changed slowly in the reciprocal shifting treatments.

Another trial was conducted with achene samples previously stored for 16 months with MC 4% at +10 and 15°C. A sub-sample was taken from each of both storage temperatures, and the MC was modified from 4 to 6% using a humid chamber with 100% RH while maintaining the same ST (+10 and 15°C) during MC modification and storage afterwards. Germination tests were conducted at 10 and 25°C.

### Sorption isotherms for “600” and estimation of equilibrium relative humidity (RH) for experimental treatments

Achenes (genotype 600) that had been stored at −18°C with MC 3.5% were used to obtain sub-samples with eight different target MC levels (≈ 4, 5, 6, 7, 8, 9, 10, 11%, DW basis), and used to build the sorption isotherms for inbred line 600. First, MC was increased to >20% in a humid chamber (RH 100%) at 20°C (about 21 d). Lower MC values were then obtained by desorption under air flow in an experimental dryer at 35°C and, to reach MC below 6%, samples were placed in a closed container supplied with dry silica. The subsamples were removed and stored as they reached the target MC (assessed gravimetrically). In this way, samples were obtained by desorption, simulating to some extent the progressive drying after harvest. The achene subsamples (20-22 grams) were placed in eight airtight 150 cm^3^ glass containers, which were fitted with a DHT22 Arduino sensor for RH and temperature on the inside of the container lid (Image S. 12 F, G) and connected to a data logger that collected data every hour. The eight containers were placed first at −18°C (freezer) and then moved sequentially to chambers set at 5, 10, 15, 20, 25 and 30°C. The containers remained 3 d at each temperature, although RH reached equilibrium within a few hours after transferring to a new temperature (SupplementaryFig. S 11 A). From these data, sorption isotherms were obtained by fitting a third-degree polynomial function for the MC data as a function of RH for each ST (see SupplementaryFig. S 11 B). These equations were used to estimate RH (and *a_w_*) for the samples used in *experiment 3*, according to their actual MC values and ST values.

Using the RH values, the water potential was also calculated according to:

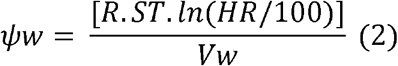

where *R* is the ideal gas constant (8.3143 J/mol.K), ST is storage temperature in K, and *Vw* is the partial molar volume of water (18.015 cm^3^mol^-1^) (Vertucci and Roos, 1993).

### Data analysis

Germination data for each storage treatment is shown as the average of 3 replicates with the standard error of mean, and GraphPad Prism 7 software was used for data presentation (Motulsky, H.G., 2003). Data for MC values (as determined gravimetrically), as well as other measurements (O_2_ level, electrical conductivity and MDA content) were analyzed by ANOVA and according to a factorial array design, with a significance level of 5% and Tukey’s test for means separation, using InfoStat software (InfoStat version 2010. Grupo InfoStat, FCA, Universidad Nacional de Córdoba, Argentina) assisted by R (R version 2.11.1 Copyright 2010. R Core Team, 2017).

### Achene relative dormancy release rate (RDRR)

Accumulated Gaussian regressions were adjusted to germination data (final germination %) obtained at different storage times using Graphpad Prism 7 software, for both 10 and 25 or 30°C incubation. The storage time required to reach 50% germination (T50) was estimated, and the output rate was calculated from its inverse (1/T50). These values were then expressed as relative to the maximum value within the same experiment to obtain the relative dormancy release rate (RDRR).

### Estimation of achene relative deterioration rate (RDetR)

The viability function of Ellis and Roberts (1980) was used to calculate the number of *p* days to obtain a viability of 50% of the population, and then the rate was obtained by calculating the inverse (1/*p*). The viability equation is:

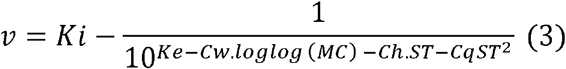

where *v* is the germination (probit) after *p* storage days, *Ki* the initial germination (probit distribution), *MC* the moisture content, *ST* the storage temperature and *Ke*, *Cw, Ch* and *Cq* species-specific constants (6.74, 4.16, 0.0329, 0.000478 respectively). Finally, the observed RDRR values and the calculated RDetR values (obtained by fitting an exponential model) were plotted against a RH and a ψw scale.

## Results

The moisture content (MC) of achenes after storage for 30 and 70 d remained close to the pre-storage values (Supplementary Table S1 and Fig. S1). Despite some fluctuations in MC data resulting in interactions between MC and ST, the overall pattern showed that MC remained stable throughout the storage period and was not consistently affected by storage temperature.

### Effects of MC and ST on germination response of achenes and embryos

Dormancy status was assessed through germination trials performed before (pre-storage) and after different storage periods. Results for one experimental year are shown in Fig. 1 (data from *experiment 2)*. Pre-storage germination scores differed among MC treatments when tested at 25°C incubation, and embryo dormancy was partially alleviated by drying achenes to MC 6 and 4% (see also Supplementary Table S2). Pre-storage embryo and achene germination scores were equally null for the four MC treatments when tested at 10°C.

**Figure 1.**
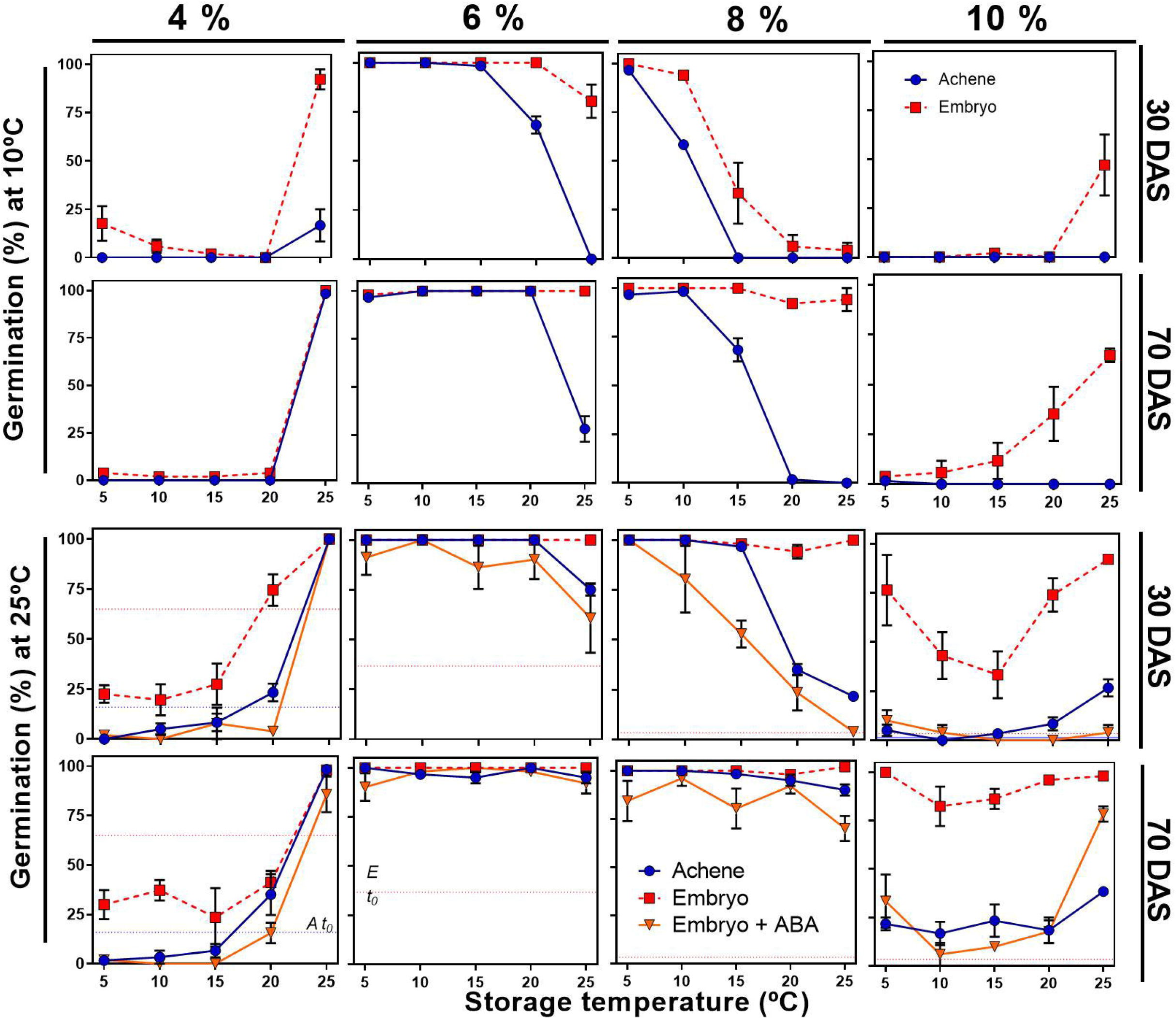
Effect of achene moisture content and temperature during storage on achene and embryo germination. Each panel shows final germination percentage of achenes and embryos as a function of storage temperature (ST: 5, 10, 15, 20 and 25°C). Germination was tested at 10°C (upper rows) and at 25°C (bottom rows) and at 30 and 70 days after storage began (30 and 70 DAS). Embryos were incubated in water and in 5 μM ABA (only at 25°C). Panels within each column correspond to a same target moisture content (MC: 4, 6, 8 and 10% dry weight basis). Data corresponds to inbred line “600” in ***experiment 2***. Germination values at the onset of storage (after adjusting achene MC to target values) are shown within each panel as horizontal dashed lines (red for embryos labeled Et_0_, and blue for achenes, labeled At_0_; when lines are not visible, germination was null). Each point represents the mean ± S.E.M. (n=3 replicate containers).

Germination response tested after 30 and 70 d of storage was deeply affected by ST and MC (Fig. 1; see also Supplementary Fig. 5 for another experimental year). Depending on achene MC, warmer storage temperatures had a promoting or an inhibitory effect on DR. For the lowest MC (4%, Fig. 1A-D), partial DR was only observed after storage at 25°C (see also Fig. 2), resulting in higher achene germination values together with increased embryo germination in water and 5 uM ABA. When stored at 20°C and below, dormancy was maintained (achene germination remained unchanged and embryo germination was even lower as compared to pre-storage values when tested at 25°C). In contrast, dormancy release was strongly promoted in achenes with MC 6% when stored between 5 and 20°C resulting in high germination values at both 25 and 10°C incubation (Fig. 1E-H). In this case, storage at 25°C delayed dormancy release resulting in lower germination values for achenes imbibed at 10°C (Fig. 1E,F) and for embryos in 5 uM ABA (Fig. 1G). This negative response to warmer ST was more pronounced in achenes with MC 8% (Fig. 1I-L) when stored between 5 and 25°C. This pattern was also followed by isolated embryos incubated in water at 10°C (30 DAS; Fig. 1I) and in 5 uM ABA at 25°C (Fig. 1K). Dormancy release was further delayed in achenes with MC 10% (Fig. 1M-P).

**Figure 2.**
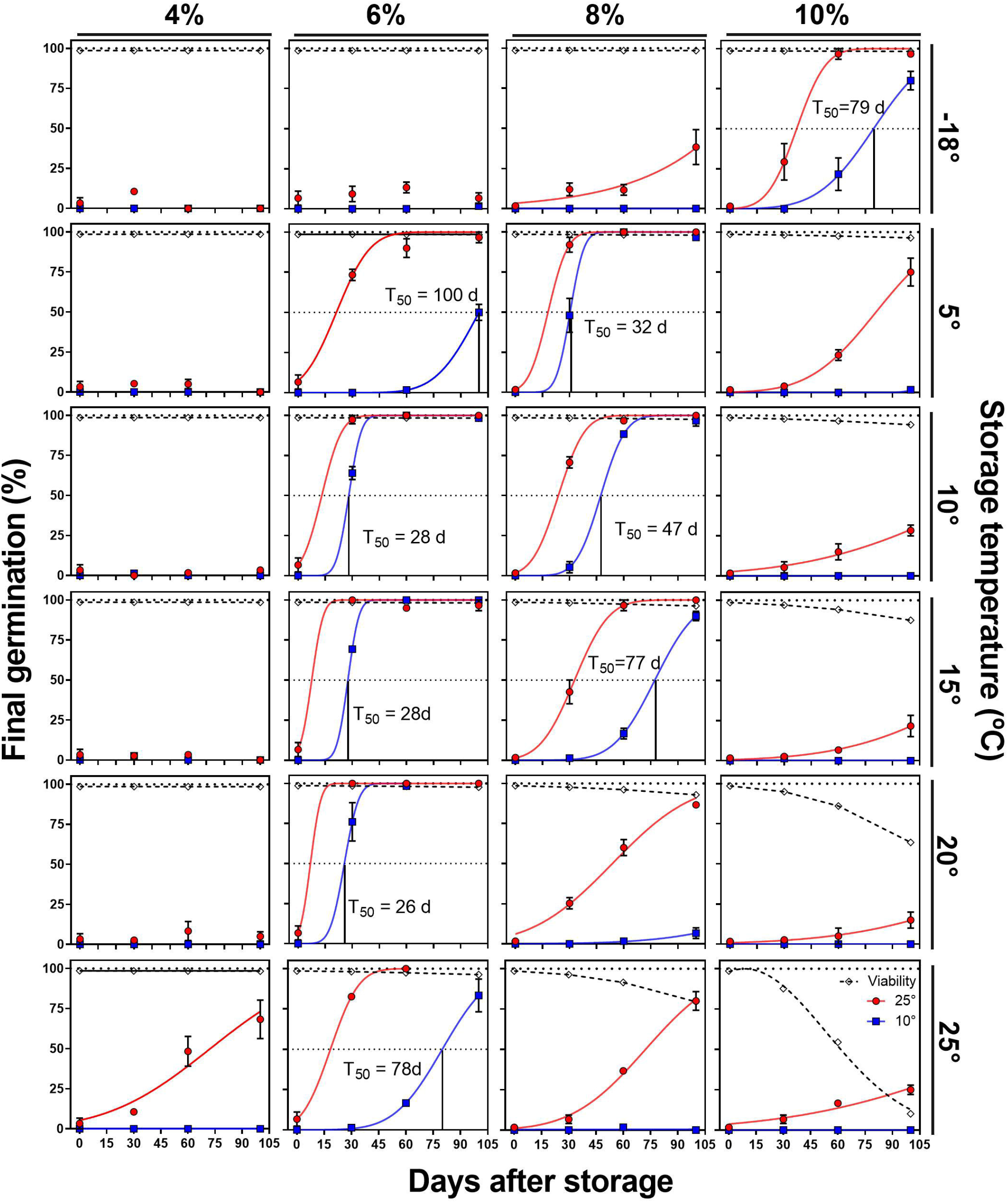
Effect of storage temperature and achene moisture content during storage on dormancy release and viability loss dynamics. Each panel shows final germination percentage of achenes at two imbibition temperatures (10 and 25°C, blue and red symbols respectively) as a function of storage time (0, 30, 70 and 100 d). Each different panel presents data for a particular MC (in columns: 4, 6, 8 and 10% dry weight basis) and storage temperature (in rows: −18, +5, 10, 15, 20, 25°C). Data belongs to genotype 600 of ***experiment 3***. Predicted viability curves are shown as dotted lines and were calculated for each storage condition using the viability function (Roberts and Ellis, 1980). Speciesspecific constants (Ke, Cw, Ch and Cq) were 6.74, 4.16, 0.0329 and 0.000478 respectively. Each point represents the mean ± S.E.M. (n=3 replicate containers).

Interestingly, a positive response to ST was again observed when combining high MC (10%) and temperatures above 15°C, where higher ST led to increased germination scores of achenes only when imbibed at 25°C (Fig. 1 O) or embryos when tested at 10 and 25°C (in water and 5 uM ABA; Fig. 1 N-P), while achene germination at 10°C remained null. A similar pattern was observed in *experiment 1* (Supplementary Fig. S 4) where achenes germinated fully when imbibed at 30°C, but failed to germinate at 10°C, even after prolonged (7 m) storage. These results support an incomplete alleviation of achene dormancy during storage with MC>8% in combination with warm (>15°C) temperatures.

Embryo germination in ABA (either when tested at 10 or 25°C) was closely related to germination of achenes in both *experiments 1 and 2*, and in both genotypes (“600” and “1579”, Supplementary Fig S2 and S3). This led to a positive and significant correlation between achene germination in water and embryo germination in 5uM ABA (p<0.0001, r=0.96, R2=0.91; Supplementary Fig. S9) for data in *experiment 2*, including all combinations of MC-ST and sampling times (30 and 70 DAS). This suggests that achene DR under different storage conditions is mediated by changes in embryo responsiveness to endogenous ABA. Possible variations in endogenous ABA in the embryo axis were also examined in *experiment 2* (Supplementary Fig. S9B). After a 70-d storage period, ABA content differed among storage treatments, with a tendency to increase towards higher ST and MC (e.g., ABA content was 551 and 717 pg.mg^-1^DW for MC4%-ST5°C and MC10%-ST25°C, respectively, different with p<0.0001). These variations in ABA content were not associated with dormancy but may reflect some degree of enzymatic hydrolysis of ABA-GE conjugates.

### Dormancy release and viability dynamics during storage

Fig. 2 shows the observed DR dynamics tested at 10 and 25°C (data from *experiment 3)* plotted together with the expected viability along a 100-d storage period. Overall, storage conditions affected dormancy release according to data shown in Fig. 1 *(experiment 2)*. In this new experiment, storage was also performed at −18°C. For achenes with MC 10%, storage at −18°C was the only condition that allowed full DR (i.e., *ca*. 85 and 100 % germination at 10 and 25°C incubation, respectively) and maintained full viability within the 100-d storage period. Achenes stored with MC 4 and 6% also maintained high viability levels at all ST during the 100-d storage period and beyond. In contrast, loss of viability within this period was predicted to occur in achenes with MC 8 and 10% in combination with ST 20°C and above. Dormancy release was also increasingly delayed by warmer ST, supporting that achenes stored under these conditions will likely lose viability before dormancy is fully released (i.e., achenes are able to germinate at 10°C).

### Effects of MC and ST on dormancy release rate followed a similar pattern across different experimental years and for two genotypes

Fig. 3 shows the relative DR rates (RDRR) obtained from germination data at 10°C incubation, for all storage treatments included in *experiments 1, 2* and *3*. Response patterns to ST under each MC condition were similar across experimental years but also when comparing two genotypes (“600” and “1579”) included in *experiment 1*. Dormancy release was inhibited by storage at low MC combined with low ST (i.e., MC 4% × ST −18 to 15°C, and MC 6% × ST −18°C). Achenes with MC 6% achieved high RDRR for ST between 5 and 25°C, with an optimal around 15-20°C above which the RDRR decreased. This negative response of RDRR to increasing ST was even more pronounced for achenes stored with MC 8% and MC 10% (for ST between −18°C and +10°C).

**Figure 3.**
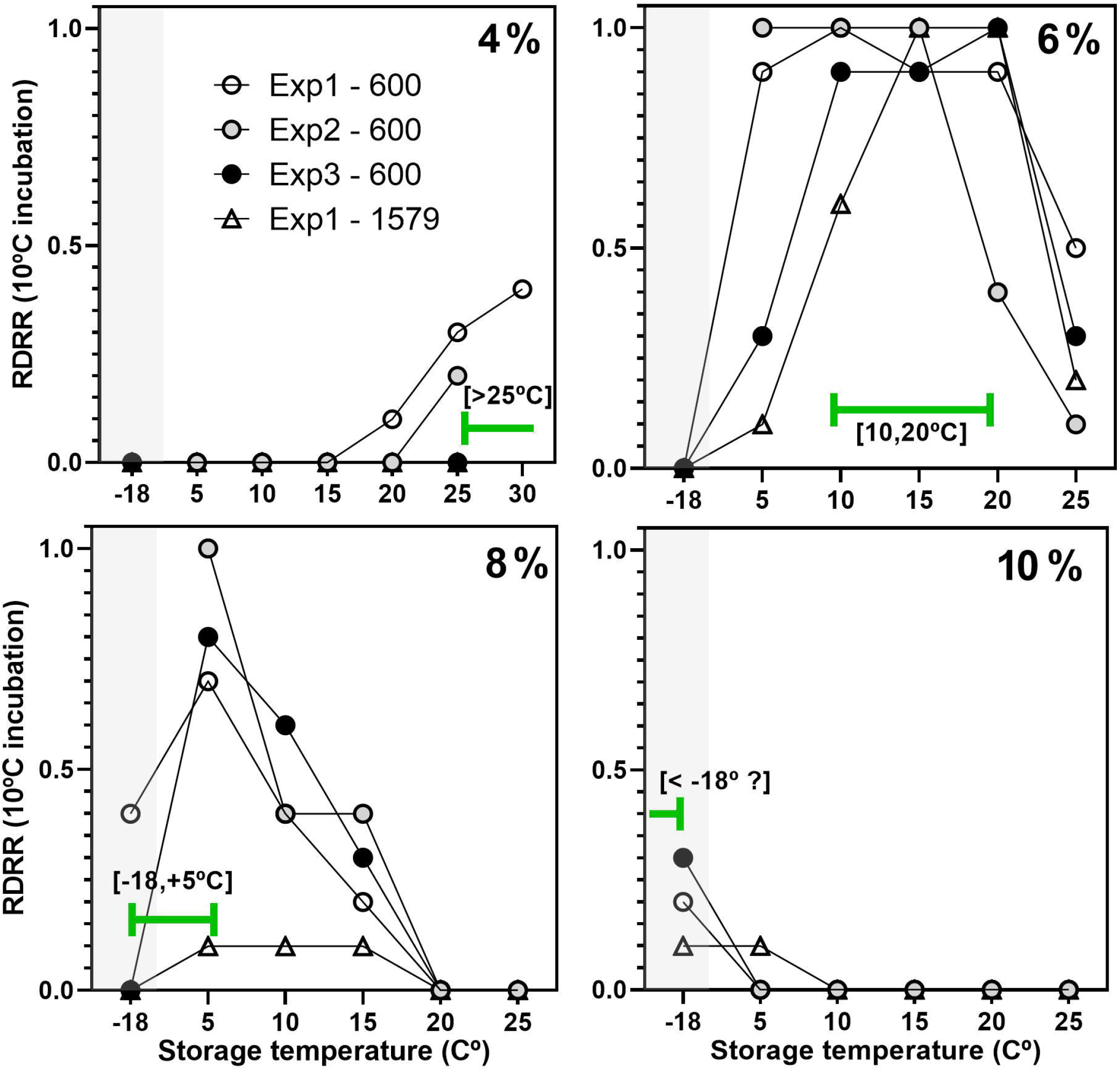
Effect of the storage environment on dormancy release velocity tested in three experimental years. Each panel shows achene relative dormancy release rate (RDRR) as a function of storage temperature (ranging from −18°C to +25 or 30°C) for achenes stored with a similar target moisture content (separate panels, MC: 4, 6, 8 or 10%, dry weight basis). Data is shown for inbred lines 600 or 1579 in different experimental years (experiments 1, 2, 3). Grey shade in each panel highlights sub-zero storage temperature. Each data point represents the estimated value of dormancy release rate (RDRR = 1/T50) for incubation temperature 10°C. Dormancy release rates are expressed as relative to the maximum value within the experimental year and genotype. Time to 50% germination (T50) values were interpolated using the fitted equations to the final germination data as shown in Figure 2 (for experiment 2) and data from experiments 1 and 3 (Supplementary Fig. S6 - 8).

### Early ageing indicators and germination response after 7 months of storage

Low germination could result from both decreased seed vigor and/or the presence of dormancy, so these processes should be distinguished carefully. The negative response to ST observed at higher MC (8 and 10%) could result from a delayed DR, or a decrease in seed vigor (anticipating loss of viability) of already non-dormant achenes. To understand better how deterioration and DR took place within our storage treatments, germination was tested in achenes from *experiment 1* after a 7-month storage period. Achene germination was tested at 10°C incubation (Fig. 4A) where dormancy is more clearly expressed. Treatments with low MC and ST (MC 4% stored at 15°C or below, and MC 6% stored at −18°C) remained deeply dormant (the isolated embryos did not germinate and did not respond to Etephon, an ethylene donor, or Fluridone, an ABA synthesis inhibitor) and remained alive after one month incubation at 10°C; Supplementary Fig. S12). Low germination of achenes stored with higher MC (8 and 10%) was reverted by Etephon (MC 8% × ST 20-25°C, and MC 10% × ST 5-10°C) suggesting a shallow-dormant state. Germinated achenes produced normal seedlings (Supplementary Fig. S12). The fresh weight of seedlings was similar across all storage treatments, including those that required Etephon to germinate (Fig. 4C). Germination scores continued to increase between 70 and 200 d (Supplementary Fig. S4) supporting that DR proceeded at a slow rate and that, within this 7-month storage period, germinative vigor as well as post-germinative growth of achenes with MC 8% were not compromised. Loss of viability was evident for achenes with MC 10% after 7-m storage at 20-25°C; most achenes were not able to germinate at 10°C and, those that did, produced abnormal seedlings (Supplementary Fig. S12F,G).

**Figure 4.**
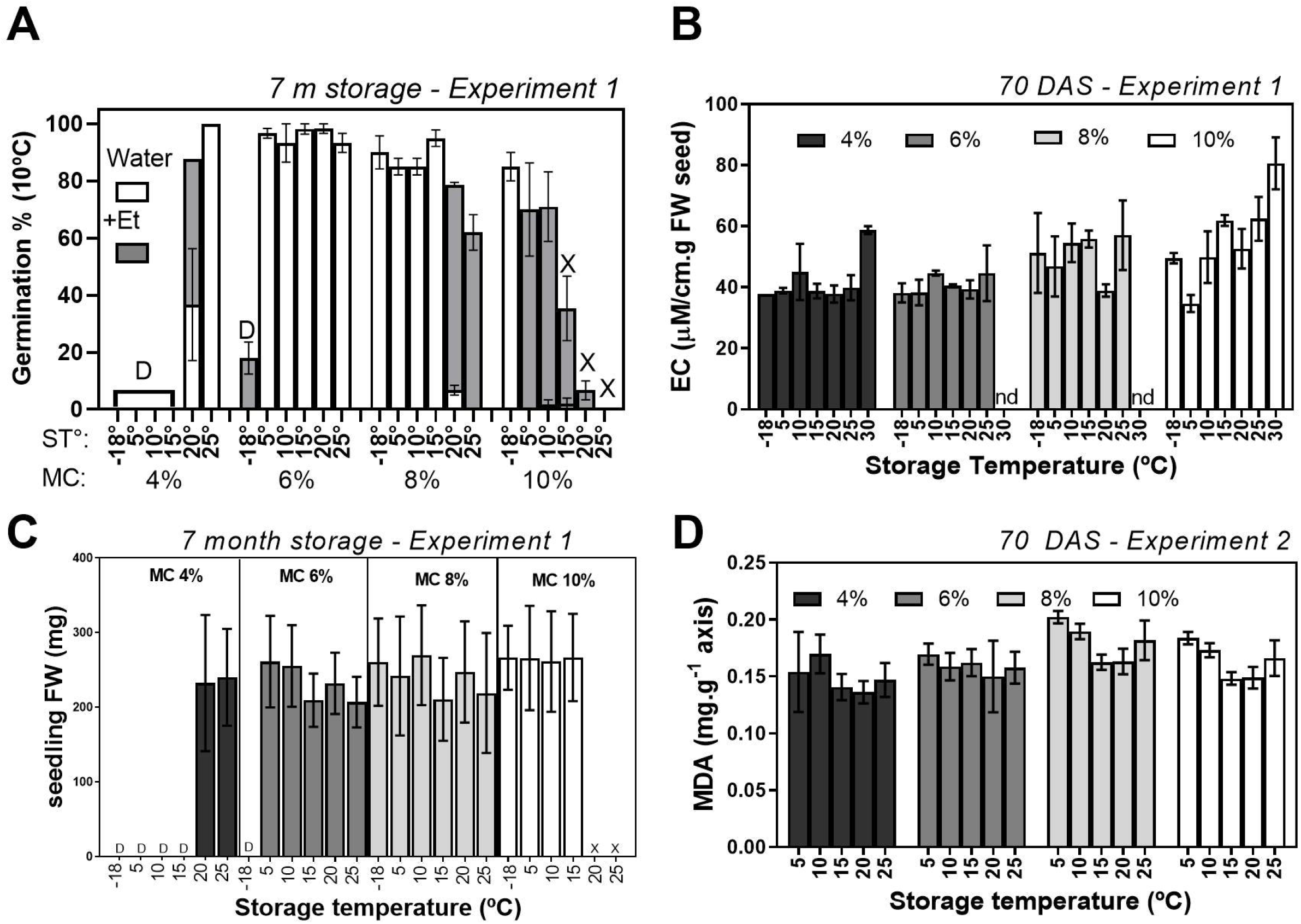
Indicators of vigor and deterioration for achenes stored under different MCxST conditions. Achene/seedling vigor was assessed after 7 months of storage through (A) germination at 10°C in water (for 15 d, white bars), or after the addition of etephon (an ethylene donor; grey bars), and (C) fresh weight of 4-d old seedlings. Data shown in (A) and (C) belong to the same germination trial with material from experiment 1 (inbred line 600). The electrical conductivity test (B) was performed with seeds (no pericarp) after 70 d of storage (Experiment 1). (D): Lipid peroxidation was measured by TBARS (expressed as MDA) in embryo axes from achenes stored for 70 d (experiment 2). In both (A) and (C), letter “D” indicates absence of germination due to deep embryo dormancy, while “X” indicates loss of viability or abnormal germination (Figure S. 13) as determined visually. Letters “nd” in (B), not determined. All data points shown are the average of 3 biological replicates (replicate storage containers; A, B, D), or n (ca. 20-25) individual seedlings (C). All error bars are SD of mean.

Solute leakage (measured by EC; Fig. 4B) is a measure of cell membrane integrity, and typically increases with ageing. In the case of sunflower seeds, solutes leak directly from (or through) the endosperm layer that surrounds the embryo. Overall, increasing MC and ST resulted in higher EC values although interaction between factors was significant (p<0.01**). Achenes with high MC (10%) were more sensitive to increasing ST (highest EC values were observed for MC 10% at 30°C). For achenes stored with lower MC (4-6%) EC values were relatively stable across ST up to 25°C (EC increased sharply after storage at 30°C for MC 4%). Lipid peroxidation in the embryo axis (assessed as MDA levels after 70-d storage period, data from *experiment 2)* did not respond clearly to MC and ST except for a slight increase in MC 8% (only MC 4% vs MC 8% differed significantly, p<0.01**; interaction NS). Overall, these results support that storage treatments most efficiently promoting dormancy release do not appear to be related to increased deteriorative/oxidative reactions that would also lead to increased EC or MDA levels. Indicators of seed coat/embryo deterioration were observed to increase mostly in treatments that delayed dormancy release (MC 8 and 10%).

### Effect of oxygen availability on dormancy release

Changes in O_2_ level during storage may reflect basal (incomplete) metabolic activity of the embryo or also non-enzymatic oxidative reactions. The O_2_ level in the headspace was measured in the closed containers prior to opening them before each germination assay in *experiment 1* (Fig. 5A). Oxygen level remained unchanged for all achene MC when stored at 15°C or below. At ST > 15°C, O_2_ levels decreased slightly (about 10% at 70-77 DAS) for MC 4, 6 and 8%, and this was similar for both genotypes. However, also in both genotypes, O_2_ level decreased markedly for MC 10% and ST 20-25°C. After 70 d, O_2_ was almost completely depleted by achenes with MC 10% stored at 25°C (inbred line 600). Since achenes were not treated with fungicides or antibiotics, the contribution of microorganisms in the pericarp to this O_2_ depletion cannot be ruled out (Domínguez *et al*., 2019). However, it is assumed that this contribution was not a main determinant of O_2_ consumption because no proliferation of fungi or other microorganisms was observed after storage.

**Figure 5.**
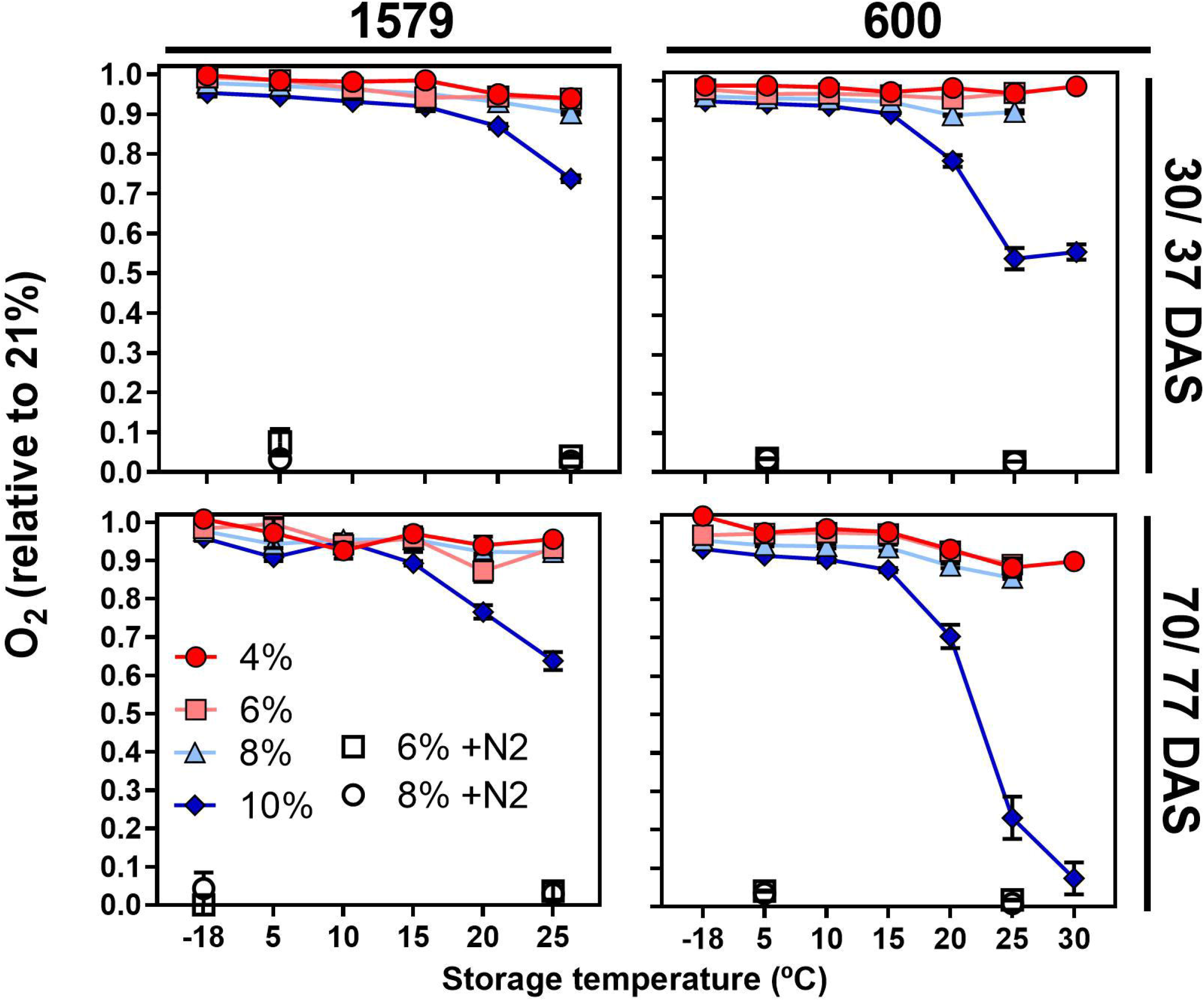
Oxygen uptake by achenes during storage under storage conditions. Each panel shows the oxygen content (expressed as relative to ambient air, 21%) in the headspace of hermetically sealed flasks (100 cm3) containing achenes (10 g) with different MC (4, 6, 8 and 10%) and stored at different temperatures. The determinations were made on two genotypes (left panels: 600 and right panels, 1579) of Experiment 1, at two moments (days after storage, DAS) during storage (upper panels: 30-37 DAS, and lower panels, 70-77 DAS). The empty symbols represent the anoxia treatments (6%, squares; 8%, circles). All measurements were performed at 25°C. Interactions between factors were significant (<0.0001***) for both times and genotypes. Each point represents the average ± SEM (n=3 replicate containers).

The effect of O_2_ availability during storage on dormancy release was also investigated in *experiments 1* and *2* by performing storage treatments under gaseous N2 (anoxia). Results shown in Fig. 6 support that the presence or absence of O_2_ in the storage containers had no detectable impact on dormancy release. Germination differed between normoxia/anoxia conditions only for achenes with MC 10%, and achene germination was higher after storage in N_2_ as compared to normoxia. Germination of isolated embryos in water or 5 uM ABA was not affected, supporting no changes in embryo dormancy status. This negative effect of O_2_ on achene germination without any apparent effect in embryo dormancy suggests the occurrence of oxidative damage to the seed coat (particularly, the living endosperm) likely affecting its role in pericarp opening during normal germination.

**Figure 6.**
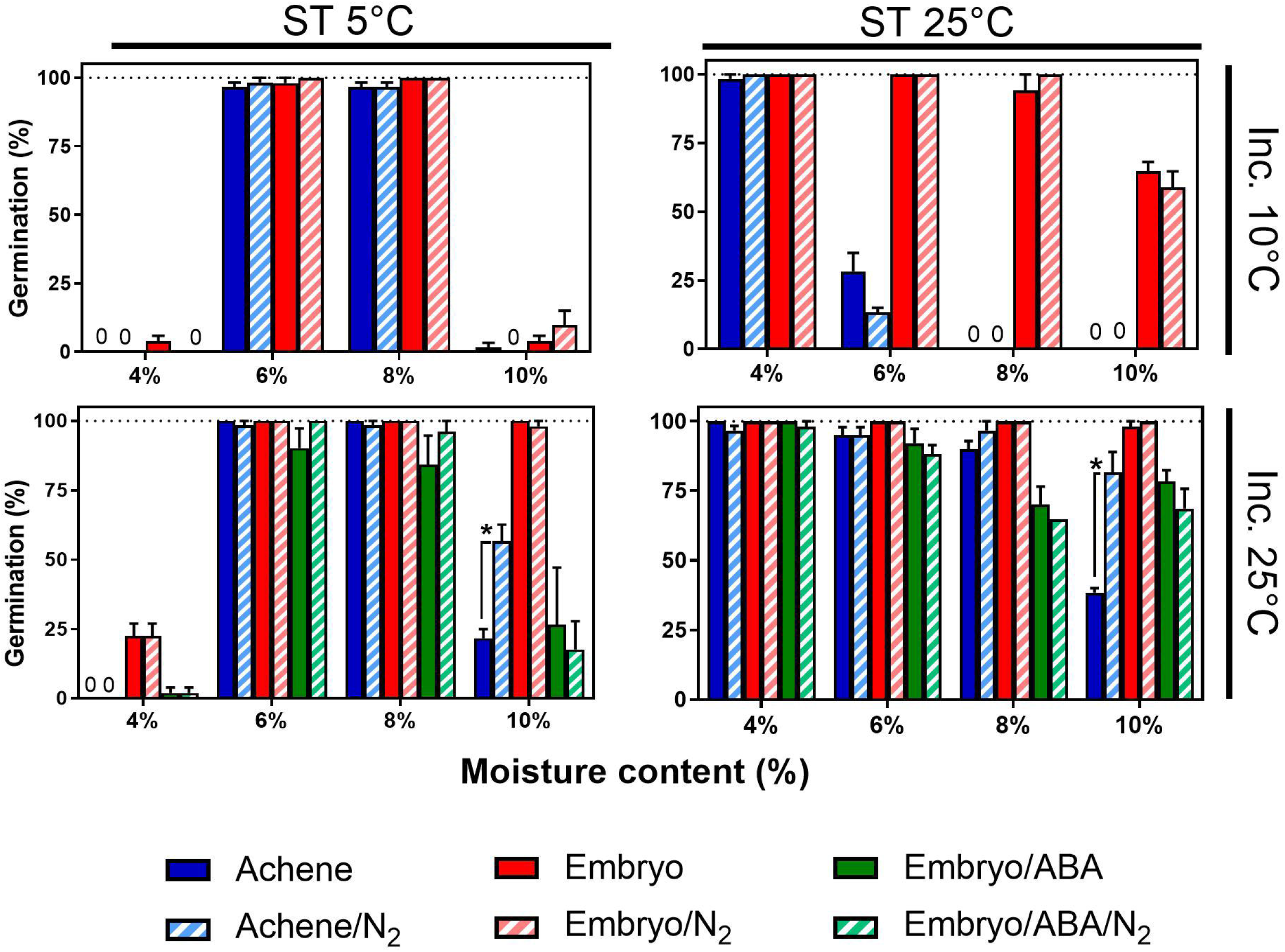
Effect of oxygen availability on dormancy release of achenes stored under different MC × ST conditions. Final germination data is shown after storage treatments combining two storage temperatures (ST 5 and 25°C, left and right panels) with four MC (4, 6, 8, 10%, nested within each panel), and tested at two incubation temperatures (10 and 25°C, upper and lower panels). Each of these MC×ST° treatments was also combined with normoxia (ambient air) or anoxia (N2). Within each panel, final germination is shown for achenes (blue bars), or embryos incubated with 5μM ABA (green bars) or in water (red bars). Results from control (air) are shown as solid bars and those from anoxia (N2) as striped bars. Data from experiment 2 (70 DAS). Effect of anoxia vs normoxia was only significant when MC 10% (shown with asterisks). Each bar represents the mean ± SE (n=3 replicate containers).

### Dormancy release rate varied along a relative humidity gradient

The RH (or *a_w_*) for each storage treatment in *experiments 1, 2* and *3* was calculated using the corresponding sorption isotherms obtained for inbred line “600” (Fig. 7 A). The RDRR values (germination tested at 10°C) for “600” in *experiments 1, 2* and *3* were plotted as a function of RH (Fig. 7C) and ψw of achenes (Supplementary Fig. S 11D). Isothermal curves were fitted to relative deterioration rate (RDetR) values predicted by the longevity model (Roberts and Ellis, 1980) and plotted along the RH scale (Fig. 7B). Four regions could be distinguished according to variations of RDRR and RDetR along the RH gradient. Region “I”, located at RH 20-40% (ψ_w_ −170 to −130 MPa), stood out for the absence of dormancy release and extremely low deterioration. This region included treatments with low MC (4%) × ST −18° to +15°C, and MC 6% stored at −18°C. In region “II”, relative RDRR increased sharply with RH between 40 - 60% (ψ_w_ −130 to −70 MPa). Storage treatments included here were MC 4% × ST 5°C, MC 6% × ST 5 −20°C and MC 8% × ST −18°C. In region “III”, located at RH between 60-85% (ψw −70 to −20 MPa), RDRR dropped with increasing RH. Included here were achenes with MC 6% stored at 25°C as well as those with higher MC and low ST (MC 8% × ST 5 - 25°C; and MC 10% × ST −18, +5°C). Within this region a sharp increase of the RDetR was predicted to occur at temperatures above 20°C. The last region (IV) (RH >85%; ψ_w_ > −20 MPa) was characterized by very low or null RDRR (assessed at 10°C) and an even sharper increase in RDetR at warm temperatures. Storage treatments within this zone (MC 10% × ST 10-25°C) also exhibited oxygen consumption and loss of vigor after 7 months, as well as an early (70 d) increase in EC (but not in MDA). The incomplete DR observed for achenes stored with high MCxST is evident within region IV as a sharp increase of RDRR tested at 25°C.

**Figure 7.**
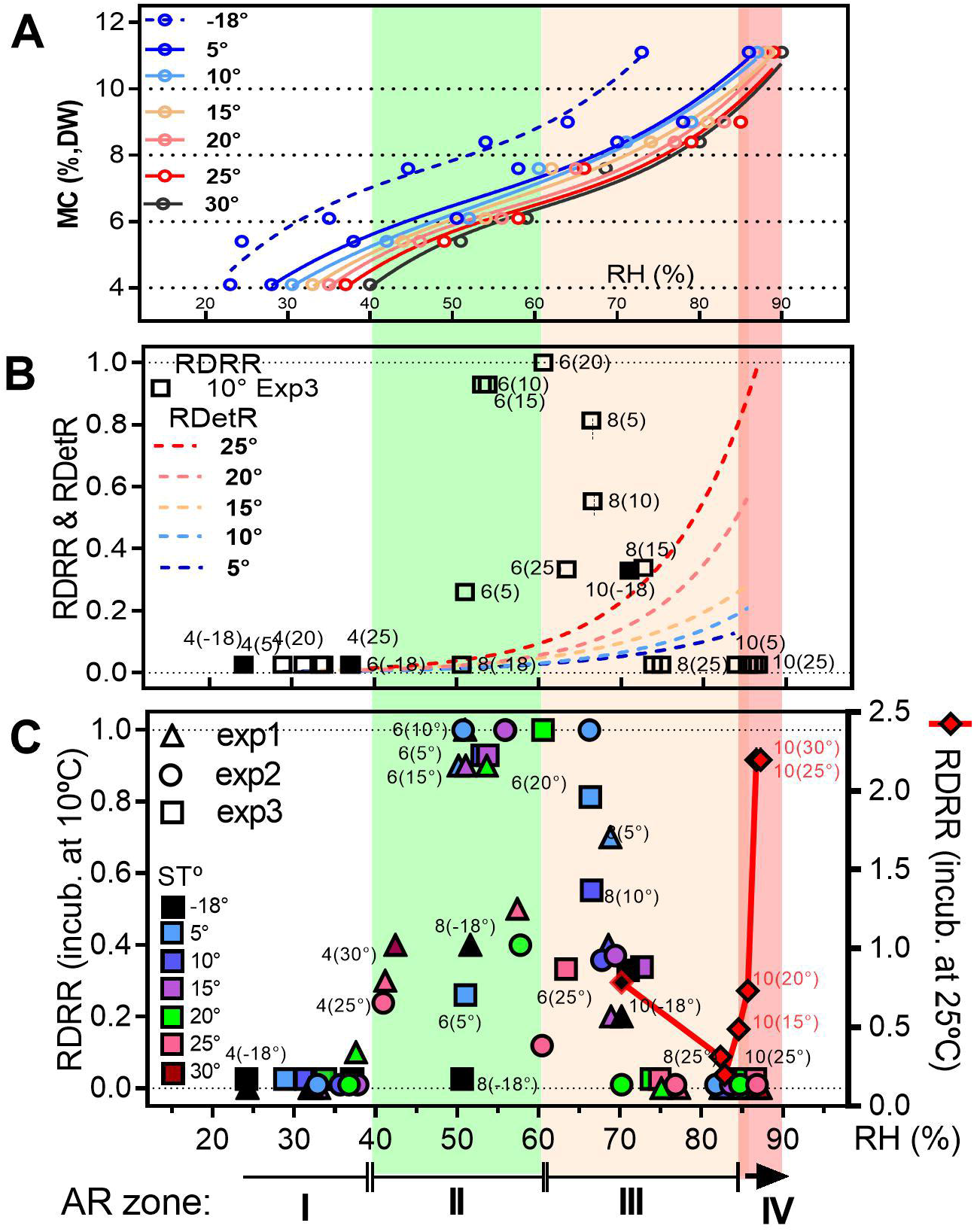
Relative dormancy release and deterioration rates as a function of RH. (A) Sorption isotherms for achenes of sunflower inbred line 600 as a function of RH. (B) Relative dormancy release rates (RDRR) for different storage treatments (MC 4, 6, 8, 10% × ST −18, +5, 10, 15, 20, 25°C; data from experiment 3, germination at 10°C) and relative deterioration rates (RDetR; calculated for ST from 5 to 25°C) plotted as a function of RH. Isothermal curves for RDetR along RH were obtained by applying the longevity equation to varying MC for fixed temperatures (5, 10, 15, 20 and 25°C). (C) RDRR (based on germination data at 10°C) for different storage treatments including data from three experimental years (represented by triangles, circles, and squares). Relative DRR for germination data at 25°C is also shown for MC 10% achenes in experiment 2 (red symbols and line; right y axis). Storage temperatures are identified with different colors (legend in panel C). Horizontal brackets show the range of RH for a single target MC (except for ST - 18°C in experiments 1 and 3; these are shown as black solid symbols). Four regions (I – IV, shaded in different colors) where identified along the RH gradient according to physiological status of achenes (see main text). Within region “I” dormancy release is inhibited, and deterioration rate is low (or beyond prediction by the longevity model); in region “II”, RDRR increases with RH (resulting from higher MC or ST), while RDetR increases slightly with a small impact of ST. In region III, RDRR is delayed as RH increases (either by higher ST or MC), while RDetR increases with a strong response to storage temperature. Region IV is associated to inhibition of dormancy release, and high deterioration rates related to metabolic imbalance and oxidative stress (mild – fast aging conditions) with the possibility of incomplete dormancy release enhanced by warmer ST.

### Moisture content variations during storage modulate dormancy release dynamics

The following experiments were aimed to investigate if dormancy release rate responds dynamically to changes in MC, particularly when shifting between regions I, II, III and IV. We expected to further discriminate between ageing and dormancy release processes and test if the dormancy status can be reverted to a deeper state (Fig. 8). Three samples of achenes with initial MC 6, 8 and 10 % were stored at 20°C. Germination tested after a 28-d storage period confirmed that dormancy release was fastest for achenes stored with MC 6% (i.e., 100 and 25% germination at 25 and 10°C incubation, respectively; Fig. 8A,B) and was delayed in achenes stored with MC 8 and 10% (germination reached 12 and 4% for achenes with MC 8 and 10% incubated at 25°C, and was null at 10°C). Lowering MC from initial 8 or 10% to a final MC 6% accelerated dormancy release, with a significant increase in germination % when tested at 25°C nine days after drying (i.e., germination increased from 24 to 78% for achenes dried from MC 8 to 6%; and from 4% to 70% for achenes dried from MC 10 to 6%). A higher dormancy release rate was also evident when tested at 10°C incubation. Germination increased from 0 to *ca* 80% within a 20-d period (between days 39 and 60 after beginning of storage), while achenes with a constant MC 8 and 10% did not exceed 10% germination even after 100 d of storage.

**Figure 8.**
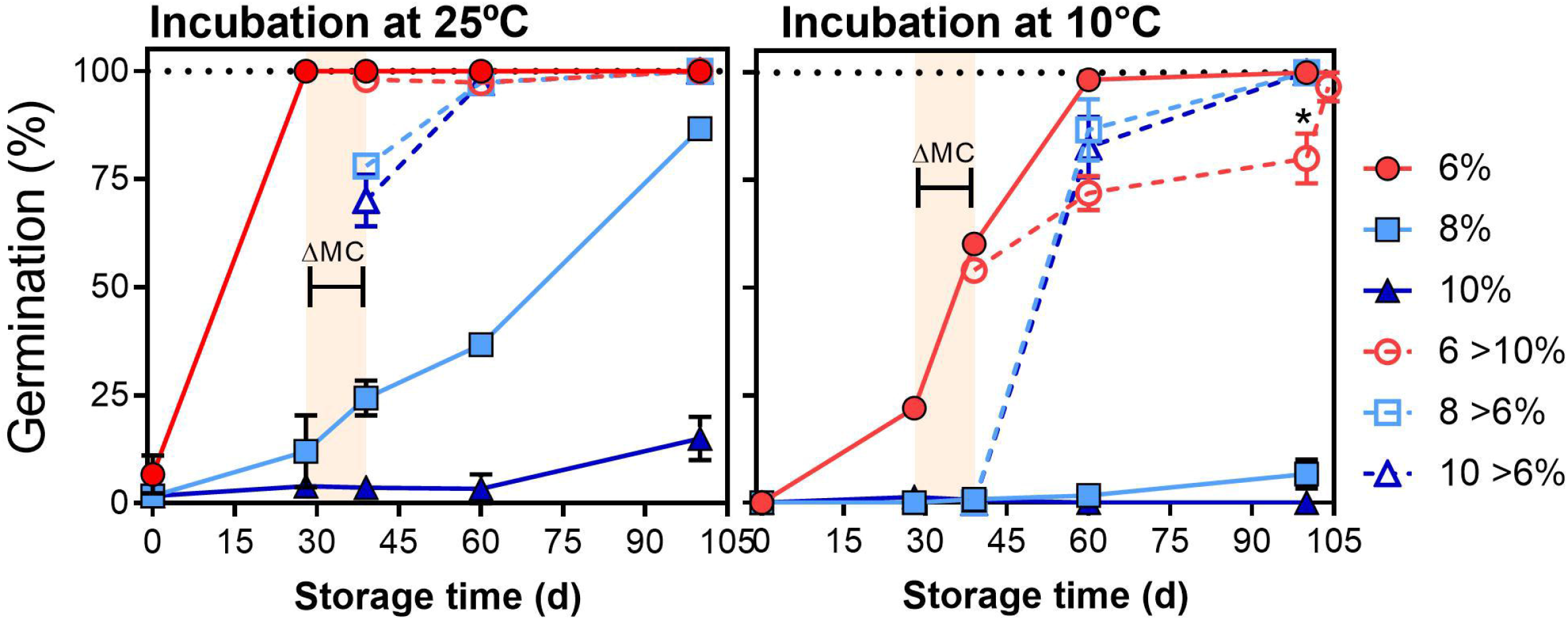
Effect of changes in MC during achene storage on dormancy release. Final germination percent (tested at 25 and 10°C, left and right panels) after different times of storage (0, 28, 38, 60 and 100 days) for achenes with different MC levels which were kept constant (6, 8 or 10% dry weight basis) or were modified during storage. After an initial storage of 28 d, achene samples were shifted to a higher (i.e., from 6 to 10%; red empty circle, dashed line) or to a lower MC (i.e., 8 to 6%, or 10 to 6%; squares and triangles, dashed lines). Shifting of MC (shown as a shaded region in each panel) took up to 10 d in a humid chamber (100% RH to increase MC from 6**→**10%) or few hours under air flow (for 8**→**6% and 10**→**6%). Storage temperature was always 20°C as well as temperature during MC manipulation. Changes in dormancy release rate can be inferred from the slope of consecutive germination scores.

Increasing MC from 6 to 10% delayed DR as compared to achenes with MC 6%. This was observed at 10°C incubation, and germination did not exceed 80% even after 100 d. At 25°C incubation, germination was 100% as before increasing MC, indicating there was no induction of secondary dormancy or loss of vigor compromising germination during this period.

To test whether fruits with very low MC that failed to germinate even after prolonged storage were still able to after-ripen, a new trial was conducted. Achenes previously stored for 16 months with MC 4% at ST 10 and 15°C (conditions preventing dormancy release) were used. A sub-sample of each of both ST was used to modify its MC from 4 to 6% while maintaining the same ST as previous. After a 40-d storage period (Table 1), samples with MC 6% were non-dormant (i.e., germination reached 100% at both 10 and 25°C incubation, and for both ST), while samples stored with 4% MC, remained absolutely dormant (0 % germination at any incubation temperature).

These results indicate that achenes stored under conditions that are inhibitory for dormancy release (e.g., high MC ≈8-10% and ST 20°C, or low 4% and < 15°C), are still able to lose dormancy at a high rate if they are dried or moistened to MC levels optimal for this process (≈6%). For samples stored with high MC, promotion of dormancy release by drying to MC 6% would be effective within a more limited time range, as deterioration proceeds at higher rates. Dormancy release is therefore a gradual process, the rate of which can be modulated according to variations in MC and ST. The results presented here support that dormancy release (at least within the range of conditions explored) occurs unidirectionally, and that modifying MC within a narrow range (4% > MC < 8%) causes important changes in dormancy release velocity. No evidence of induction into secondary dormancy was observed.

## Discussion

It has been recognized for long that changes in viability and primary dormancy occurring in orthodox “dry” seeds are greatly influenced by variations in moisture and temperature. Nevertheless, few studies considered complete factorial designs wide enough to explore MCxST interactions on DR (see Supplementary Table S3). For example, Foley (1994) reported an inverse relationship between optimal ST and MC for dormancy release of *Avena fatua* cariopses, while the opposite pattern was reported for sunflower and Arabidopsis by Bazin *et al*., (2011a) and Bassbouss-Serhal *et al*. (2016). Apparent differences between our study and results by Bazin *et al*. (2011a) may result from different experimental approaches and the ranges of MC and ST explored. Bazin *et al*., focused on embryo dormancy (only naked seeds were tested) and DR was enhanced by low ST when MC was below 4%, and by high ST when MC was high, >10%. In our study, the very low MC range (<4%) was not included. On the other hand, the positive response to ST at high MC (10%) observed by Bazin *et al*. is consistent with “incomplete” dormancy release of achenes reported here (allowing achene germination only at high incubation temperature). These storage conditions belong to region IV in our model for relative RDRR vs RH (Fig. 7), where DR is faulty or incomplete (although embryo dormancy decreases, achenes don’t germinate at 10°C) and deterioration rates are comparatively high. Dormancy release under these high MC and ST conditions may involve oxidative reactions as it coincides with progressive oxygen consumption (Fig. 5), higher EC values (Fig. 4) and visible abnormalities after 7 m storage (SupplementaryFig. S11B-G). Nevertheless, this partial alleviation of dormancy also took place under anoxia, suggesting that ROS from other sources may be enough to trigger these changes.

The RDRR vs RH (as a proxy for *a_w_*) model presented here assumes that reactions leading to dormancy release in “dry” achenes are limited by *a_w_*. Relative humidity below *ca* 40% inhibited DR, while increasing RH between 40 and 60% promoted this process, with a positive response to storage temperature (although the ST range depended on MC). Region II of the RDRR model covers a narrow range within the second region of the sorption isotherms and is consistent with the optimal conditions for dry after-ripening reported in the literature, i.e., within the second region of sorption isotherms, around RH 50% (Leopold *et al*. 1988; Esashi *et al*. 1993; Foley 1994; Bazin *et al*., 2011; Bassbouss-Serhal *et al*., 2016). Increasing RH from 60 to 75% by rising MC or ST (region III of our model, still within second region of sorption isotherms >5°C) resulted in a pronounced decline in RDRR.

In parallel with changes in RH (or *a_w_*), reactions in low hydrated tissues may also respond to qualitative changes in the physical status of water. As water is withdrawn and solute concentration increases, the cytoplasm enters a glassy state. Properties of the glassy matrix also depend on chemical composition of the solutes. Among the different properties describing a glass, the Tg represents the melting temperature (Leopold *et al*., 1994; Ballesteros and Walters, 2011, 2019). The Tg decreases as MC increases. Phase diagrams obtained for seeds from soybean axis and maize embryos show that Tg values decrease roughly from *ca* +50°C to −30/-60°C as MC increases from 5 to 20% (DW basis). Sorption isotherms and phase diagrams are widely used to interpret processes in the low-hydration range but are rarely combined in a single study. Although in this work we did not measure the Tg in our samples, a clue to whether region II for optimal dormancy release is related somehow to the glassy transition can be found in Ballesteros and Walters (2019) who investigated biophysical properties of embryo axis of pea and soybean in relation to ageing. Using thermal dynamic mechanical analysis (TDMA) these authors observed that fluctuations between fluid and solid states occur above and below about 50–60% RH at about 20–30°C. These conditions coincide roughly with the boundary between regions II and III as presented in our model of RDRR vs RH. This suggests that qualitative changes in DR response among both regions may reflect changes in the physical status of cellular water as it transitions from glassy −in region II- to a more fluid state along region III and towards region IV. A conceptual model is presented (Fig. 9 A) where dormancy release is related to a probable phase transition. The different regions (I-IV) described in the RDRR vs RH model are now shown as areas between cardinal temperatures (minimum, optimal, and maximum) for dormancy release, with the particularity that these are not fixed values but decrease as MC increases and run parallel to the expected Tg. Cardinal temperatures defining boundaries between areas I-IV are based on the observation of heat maps for RDRR, RH and oxygen consumed during storage (Fig. 9B-D).

**Figure 9.**
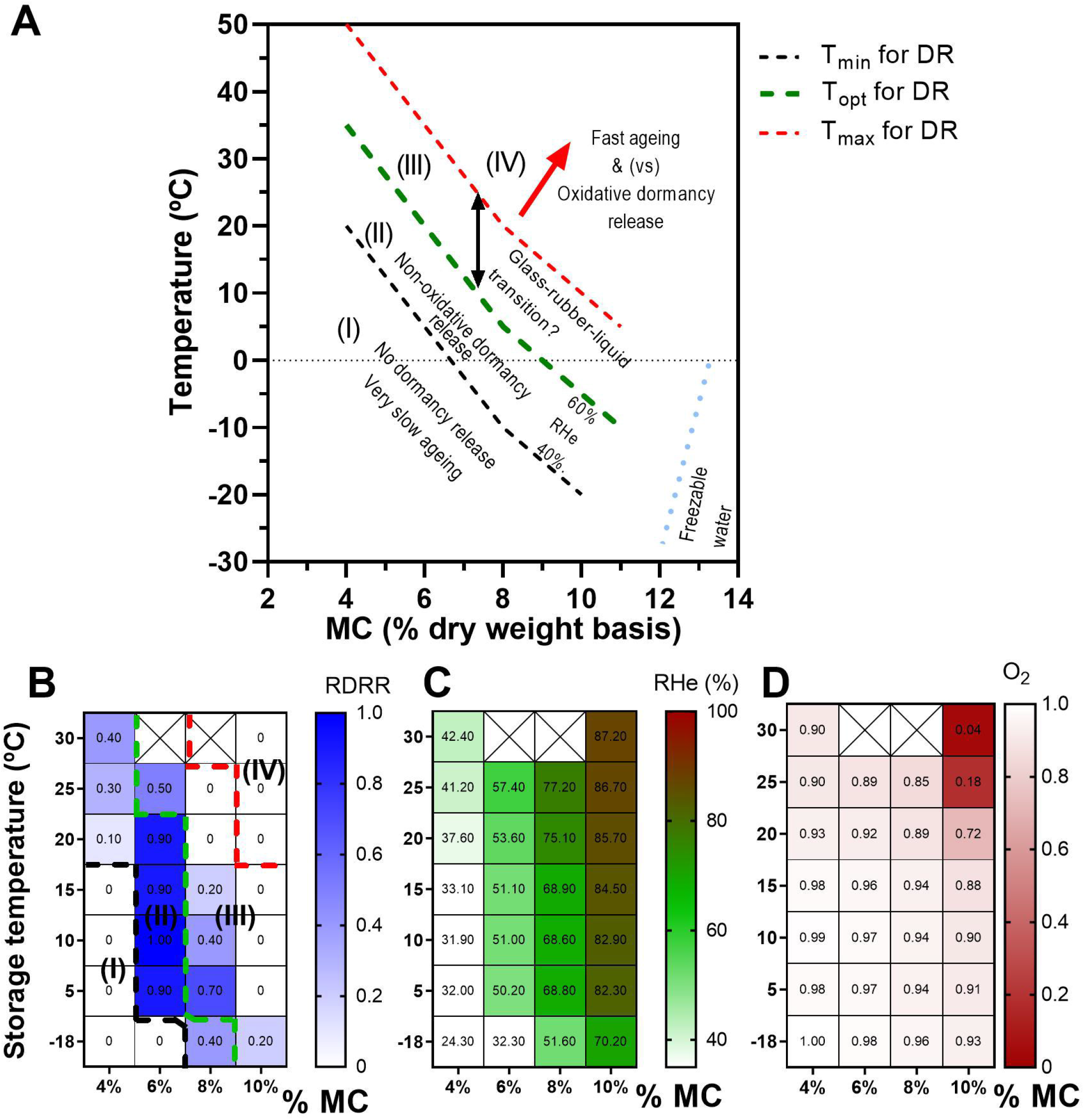
Conceptual diagram showing cardinal temperatures (minimum, optimal and maximum) for DR as a function of achene MC. (A) Regions I-IV represent different physiological states regarding dormancy release and deterioration as reflected by heat maps below for relative DR rate (B), equilibrium relative humidity (C) and oxygen consumption (D; shown as a fraction of ambient air) after 70 d of storage. In region (I), DR is null, and longevity is high. Cellular water is in the glassy state. No enzymatic reactions allowed in MLO (see main text) at very low Aw. In region (II) DR is promoted between Tmin – Topt and is non-oxidative. Ageing is very slow. Cellular water is in the glassy state. Possible enzymatic catabolic reactions in localized MLO could take place leading to decreased sensitivity to ABA resulting in DR. In region (III) DR rate slows down between Topt and Tmax. Ageing rate increases. Cellular water in glassy state transitions to non-glassy. In region (IV) DR (non-oxidative) is very slow or null, but oxidative reactions leading to partial DR and ageing increase.

Why is dormancy release favored between RH 40-60%, and slowed down or even inhibited at higher RH? A possible explanation for this pattern is that the type of reactions that allow DR at low RH (40-60%) are favored in the glassy state but become less efficient as the system transitions to non-glassy. The possibility that certain enzymatic reactions may be enhanced in the glassy state is based on an idea first suggested by Leubner-Metzger (2005) regarding the formation of “mobility pockets” and recent findings supporting localized molecular mobility within the glassy matrix. These may involve sugars and organic acids (highly abundant in seeds) which can form natural deep eutectic solvents (NaDES; Farrant and Hilhorst, 2021; Choi *et al*., 2011). Also, a non-aqueous liquid phase could be formed by condensation of intrinsically disordered proteins (IDPs; Dorone *et al*., 2021) which undergo liquid-liquid phase separation and allow the formation of membrane-less organelles (MLO) where specific biochemical reactions can proceed (Dorone *et al*., 2021; Boeynaems *et al*., 2018).

In this work we didn’t directly investigate the type of reactions involved in DR, but we do know that these reactions lead to reduced embryo responsiveness to ABA, without an obvious link with oxidative reactions. Also, results presented here support that DR is favored by combinations of MCxST that run along the Tg (Fig. 9). A close relationship between the glassy state and storage conditions promoting seed longevity was first observed by Leopold *et al*. (1994). These authors showed that variations in maximum ST for long term storage of seeds as a function of MC, overlapped with the phase diagram for the Tg. It is noteworthy that dormancy release in our study proceeded fully at sub-zero temperatures in achenes stored with MC 8-10% (in which water is structured and non-freezable). Preliminary data of Tg obtained by differential scanning calorimetry of de-fatted embryo axes of “600” supports that Tg is below 0°C for a MC in equilibrium with RH 72% at 20°C (data not shown). If this Tg value obtained for de-fatted axes is representative of intact achenes with MC *ca* 8%, where optimal temperature for DR appears to be somewhere between −18°C and +5°C (Fig. 3), then an association between MCxST conditions favoring DR and the glassy state can be hypothesized. More measurements are needed to construct the phase diagram in this genotype and confirm this hypothesis. Ballesteros and Walters (2019) investigated biophysical properties of embryo axis of pea and soybean in relation to ageing. They proposed that fluctuation between fluid and solid states occur above and below about RH 50–60% and 20–30°C and have profound effects on physiological change. Additional parameters reflecting other properties of the glassy matrix (related to glass fragility) provided by the TDMA were used to explain differences in longevity between the two species. These authors observed relaxations within the glassy matrix that may reflect localized regions with mobility. These regions or “mobility pockets” may be necessary for the reactions leading to dormancy release but also could be favored within the glassy state and dissolved in the more fluid state. It can be speculated that too much water (or temperature above Tg) removes the glassy state by disorganizing not only the glass, but also affecting (dissolving) enzymes or substrates included in an MLO formed by NaDES and/or IDPs. On the other hand, within the glassy state, too little water could inhibit enzyme activity in the NaDES. This may explain variation in RDRR observed within region II, and inhibition in region I. For example, Choi *et al*. (2011) observed that the enzymatic activity of laccase dissolved in a NaDES was shown to rely on the addition of water. So, formation of mobility pockets (or MLOs) within the glassy matrix may enhance enzymatic-catabolic-reactions involving proteins that undergo phase separation. Recently it was demonstrated that LEA6 (a LATE EMBRYOGENESIS ABUNDANT protein) undergoes phase-phase separation in desiccation tolerant *Artemia franciscana* cysts (Belott *et al*., 2020). AfrLEA6 forms MLOs that remain fluid even after total dehydration. Similar MLOs could be expected to form in dehydrating seeds involving LEA or prion-like proteins (Dorone *et al*., 2021) and NaDES. Because dormancy release in sunflower achenes appears to be governed by attenuation of ABA signaling in the embryo, catalytic post-translational modifications of key positive regulators of the ABA signal are good candidates to be investigated. For example, changes in phosphorylation status of ABI5 (which is also an IDP protein; Dorone *et al*., 2020) by phosphatases might lead to desensitization to ABA (Dai *et al*., 2013). The idea of enzymatic activity resulting in dormancy alleviation in the dry state appears to contradict the prevailing model of chemical (non-enzymatic) oxidative reactions. A role for ROS in dormancy alleviation was first proposed by Esashi *et al*., 1993, and long supported in the literature afterwards (Oracz *et al*., 2007; Morscher *et al*., 2015; Buijs *et al*., 2018; Bailly, C.; 2019). An oxidative mechanism may be involved in partial dormancy release observed in region IV together with fast ageing rates. In contrast, promotion of full DR involving loss of sensitivity to ABA (as observed within region II along the RH gradient) may involve a different, non-oxidative mechanism. Although subtle oxidative reactions cannot be discarded, we speculate that ROS formation would be low enough not to have any physiological impact within the time frame leading to dormancy release.

## abbreviations

AR: After-ripening
ST: storage temperature
MC: moisture content
DR: dormancy release
RDRR: relative dormancy release rate
RDetR: Relative deterioration rate
MLO: membrane-less organelle
NaDES: natural deep eutectic solvent
IDP: intrinsically disordered protein

## Supplementary data files

**Table S 1**. ANOVA for moisture content data (observed values) within experiments.

**Table S 2.** Pre-storage germination data for achene samples used in *experiment 2*.

**Table S 3.** Published works exploring effect of MC and temperature on dry after-ripening.

**Figure S 1.** Moisture content values for experimental treatments.

**Figure S 2.** Germination percentage of achenes and embryos (in water and ABA) as a function of storage temperature for inbred line 600 in *experiment 1*.

**Figure S 3.** Germination percentage of achenes and embryos (in water and ABA) as a function of storage temperature for inbred line 1579 in *experiment 1*.

**Figure S 4.** Temporal changes in achene germination for storage treatments applied to inbred line 600 in *experiment 1* (including prolonged storage).

**Figure S 5.** Germination percentage of achenes as a function of storage temperature for inbred line 600 in *experiment 3*.

**Figure S 6.** Dormancy release dynamics (fitted curves) for storage treatments (MCxST) in *experiment 1*, inbred line 600.

**Figure S 7.** Dormancy release dynamics (fitted curves) for storage treatments (MCxST) in *experiment 1*, inbred line 1579.

**Figure S 8.** Dormancy release dynamics for storage treatments (MCxST) in *experiment 2*, inbred line 600.

**Figure S 9.** Achene dormancy is correlated with embryo responsiveness to ABA but not with ABA content.

**Figure S 10.** Effect of oxygen (anoxia treatments) on dormancy release.

**Figure S 11.** Moisture isotherms, RH measurements, Van’t Hoff plot, relative dormancy release rate values as a function of water potential.

**Figure S 12.** Images of 7-month storage experiment to assess seed vigor and dormancy.

**Figure S 13.** Images of experimental setups.

## Acknowledgements

We are grateful to Cristian “Messi” Escudero, Maxi “la fiera” Rodríguez and Mirta “Piti” Tinaro for their valuable technical help during the experiments. This work was funded by the Universidad de Buenos Aires (research grants UBACYT No.20020170100599BA, 2018–2021), and by the National Scientific and Technological Research Council of Argentina (CONICET; research grant PIP 2015–2018, No. 11220130100669) and by the National Agency for promotion of Science and Technology (ANPCyT, PICT 2018-3546). Funding was obtained by D. Batlla (as PI) and M.V.Rodríguez. This whole work is part of the doctoral thesis of G. J. Arata, who was recipient of a doctorate fellowship from CONICET under direction of M. V. Rodríguez and Co-direction of D. Batlla.

## Author contributions

MVR, GJA and DB: Conceptualization. MVR: Supervision. PVD and GJA: Methodology; GJA: Investigation; MVR and GJA: Visualization; MVR and GJA: Writing original draft; MVR: Writing-Review and editing; MVR and DB: Funding acquisition.

## Data availability

All plant materials are publicly available as well as the original data upon request to the authors.

## Tables

**Table 1:** Effect of increasing MC of achenes after prolonged storage at 10 and 15°C with MC 4%. Achenes with MC of 4% (dry weight basis) were first stored for 16 months at 10 or 15°C. After first storage, sub-samples were taken and their MC was increased from 4 to 6%, and then stored for 40 d at the same storage temperatures as previous (10 and 15°C). After second storage (16 m + 40 d) germination of achenes was tested at two incubation temperatures (10 and 25°C). Each value indicates the mean ± S.E.M. (n=3).

| Storage<br>T (°C) | MC<br>(%) | Incubation<br>T (°C) | Germination<br>(%) |
| --- | --- | --- | --- |
| 10 | 4 | 10 | 0 ± 0 |
| | | 25 | 0 ± 0 |
| | 4→6 | 10 | 97 ± 3 |
| | | 25 | 100 ± 0 |
| 15 | 4 | 10 | 0 ± 0 |
| | | 25 | 0 ± 0 |
| | 4→6 | 10 | 100 ± 0 |
| | | 25 | 100 ± 0 |

